# A robust platform for BaEVRless-lentiviral synthesis and primary natural killer cell transduction

**DOI:** 10.1101/2024.04.03.587896

**Authors:** Yi-Jun Lan, Quoc Viet Nguyen, Tsu-Lan Chao, Kuo-Lun Yeh, Steven Lin

## Abstract

Lentiviral vectors are invaluable tools for genetic modification in human cells for research, biotechnological and clinical applications. However, certain cell types, such as primary human natural killer (NK) cells, present challenges in lentiviral transduction. Overcoming this limitation requires specific pseudotype modifications. BaEVRless-pseudotyped lentivirus (BaEVRless-LV) has shown promise in efficiently transducing human NK cells, B cells, and hematopoietic stem cells (HSCs). BaEVRless, a modified envelope protein derived from Baboon endogenous retrovirus, targets ASCT receptors in human cells. While effective for several immune cell types, BaEVRless-LV production in standard HEK293T cells is challenging. During lentiviral synthesis, BaEVRless protein induces hyper cell fusion, leading to rapid HEK293T cell death and reduced BaEVRless-LV titers. To solve this problem, we used CRISPR genome editing to knockout (KO) the *ASCT2* gene in HEK293T cells, thereby abolishing BaEVRless-induced cell fusion. Using the *ASCT2*-KO cells and an optimized viral production protocol, we efficiently packaged high titers of BaEVRless-LV encoding various transgenes, including *turbogfp*, chimeric antigen receptor (CAR), and a pooled CRISPR sgRNA library. Our robust BaEVRless-LV synthesis platform is readily adaptable for manufacturing cell therapeutics and enables advanced research techniques such as CRISPR genetic screens in primary NK cells.

## Introduction

Lentiviral vectors are powerful genetic tools to introduce and stably express transgenes in human cells. The modular nature of lentiviral packaging system simplifies virus production, enhances the safety during handling, and enables rapid exchange of the viral components and transgenes. Lentiviral tropism can also be altered by changing the pseudotype envelope protein, allowing the virus to target specific cell types (Dull et al., 1998). Despite advance in nonviral genetic modifications and delivery methods, lentiviral vectors remain indispensable for a wide range of research, biotechnological and clinical applications.

NK cells are a promising cell type for adoptive immunotherapy owing to their diverse surveillance and clearance mechanisms against malignant cells (Guillerey et al., 2016). Clinical trials have demonstrated the efficacy and safety of allogeneic NK cell transfer in treating leukemia, paving the way for off-the-shelf therapy (Liu et al., 2020). While retroviral and lentiviral transduction are standard methods for genetically modifying NK cells, the conventional vesicular stomatitis virus glycoprotein (VSV-G) pseudotyped virus is inefficient, as NK cells lack the low-density lipoprotein receptor (LDLR) necessary for VSV-G binding (Sanctis et al., 1996). Gamma retroviruses, such as RD-114 and Baboon endogenous retrovirus (BaEV), have shown greater efficiency in transducing NK cells (Colamartino et al., 2019). Incorporating the envelope proteins of these retroviruses into lentiviral vectors has been shown to enhance NK cell transduction, highlighting their potential in adoptive immunotherapy (Colamartino et al., 2019; Imai et al., 2005).

Virology studies have revealed that BaEV envelope protein contains a 16-amino acid R-peptide in its cytoplasmic domain, which acts as a negative regulator of viral fusogenicity (Lucas et al., 2010). A modified variant, BaEVRless, lacking the fusion inhibitory R-peptide, exhibits significantly enhanced transduction efficiency in NK cells, B cells, and HSCs (Girard-Gagnepain et al., 2014). However, the synthesis of BaEVRless-LV poses challenges. BaEVRless induces cell fusion, leading to syncytia formation, HEK293T cell death, and reduced viral titers (Noguchi et al., 2023; Girard-Gagnepain et al., 2014; Bauler et al., 2020). This issue not only compromises BaEVRless-LV quality and increases production costs but also hampers BaEVRless-LV-based research applications. Currently, the only available solution is the poly-L-lysine coating of culture plates to enhance cell adhesion; however, this approach does not prevent syncytia formation (Noguchi et al., 2023).

In this study, we used CRISPR genome editing to modify HEK293T cells, aiming to eliminate BaEVRless-induced cell fusion. Our approach was guided by previous findings indicating that the sodium-dependent neutral amino acid transporters ASCT1 and ASCT2 serve as receptors for BaEVRless (Tailor et al., 1999; Rasko et al., 1999; Marin et al., 2000). Using Cas9 ribonucleoproteins (Cas9 RNP), we successfully KO the *ASCT1* and *ASCT2* genes in HEK293T cells and determined that ASCT2 plays a primary role in mediating BaEVRless-induced cell fusion.

The *ASCT2* KO cells exhibited normal growth characteristics and could be maintained and transfected with high efficiency, similar to the parental HEK293T cells. Importantly, the *ASCT2* KO cells seamlessly integrated into the standard adherent cell-based lentiviral packaging system, requiring no modifications to the cell culture or transfection procedures.

Using the *ASCT2* KO cells and an optimized viral production protocol, we achieved a remarkable 50-fold increase in BaEVRless-LV titer for various transgenes, including *turbogfp*, CAR and a pooled CRISPR sgRNA library. Using our platform, we efficiently transduced primary NK cells to generate CAR-NK cells. Furthermore, we provided the first demonstration of CRISPR KO screen in primary human NK cells and identified kinase genes crucial to NK cell proliferation. Our BaEVRless-LV synthesis platform can accelerate the development of immune cell therapeutics and enable large-scale lentiviral-based research techniques, such as CRISPR genetic screens.

## Results and Discussion

### BaEVRless induces cell fusion and disrupts lentiviral synthesis

To understand the challenge posed by BaEVRless, we shall briefly review the synthesis of lentiviral vectors in HEK293T and related cell lines. The process begins with the transfection of lentiviral packaging plasmids containing genes for Gag, Tat, Pol, Rev, and pseudotype envelope proteins, along with a transfer plasmid encoding the desired transgene (Figure 1A). Expression of these viral proteins and lentiviral RNA transcript initiates the assembly of lentiviruses. These viruses are then secreted from the HEK293T cells into the culture medium, which is subsequently harvested for viral purification.

**Figure 1.**
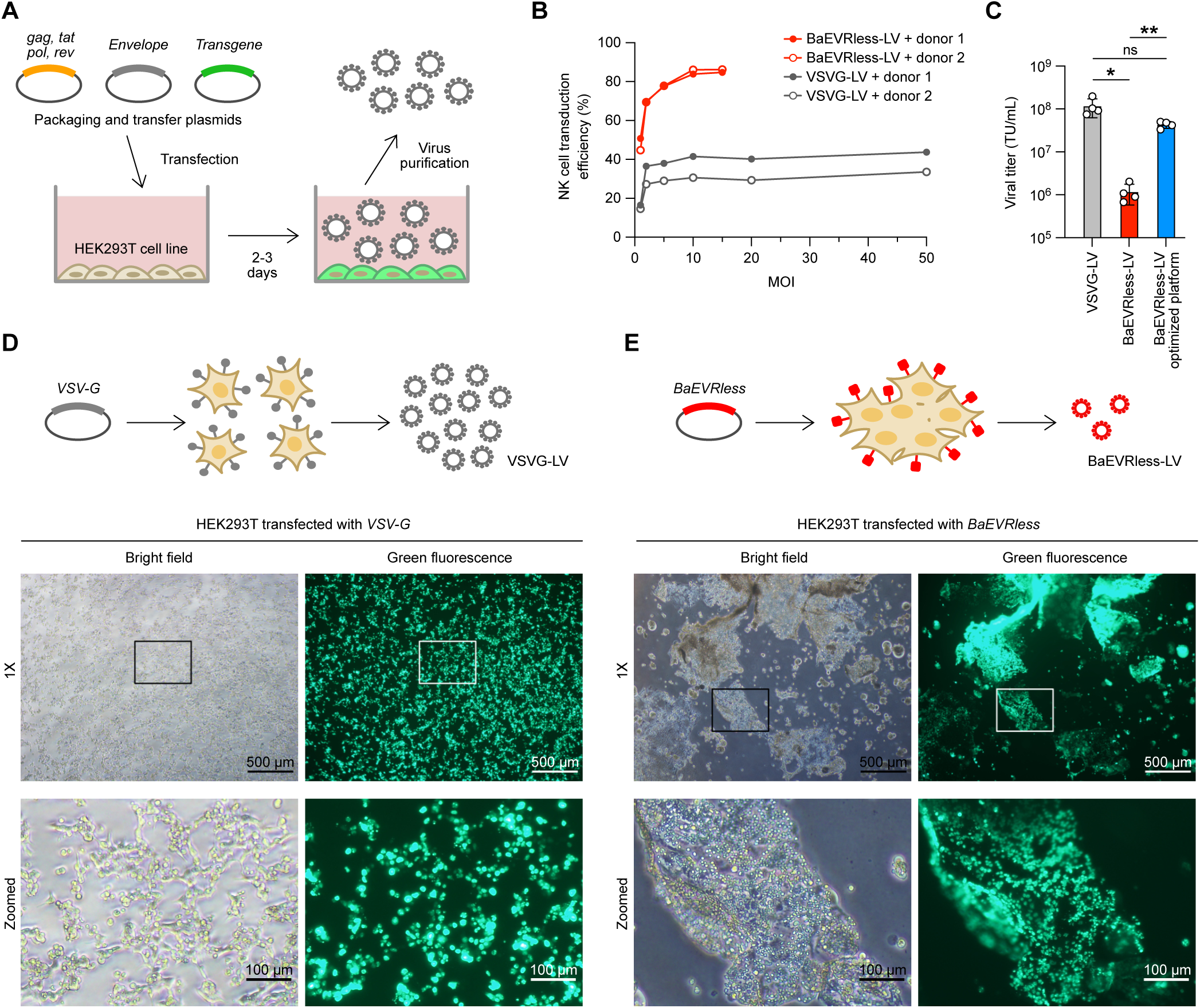
BaEVRless induced cell fusion during lentiviral production. **(A)** Overview of lentiviral synthesis by transfection of packaging and transfer plasmids in HEK293T cells. The lentiviral tropism is determined by the envelope gene. **(B)** Transduction efficiency of primary NK cells from two donors using BaEVRless versus VSV-G pseudotyped lentiviruses (BaEVRless-LV vs. VSVG-LV) at different multiplicities of infection (MOI), determined by turboGFP expression using flow cytometry. **(C)** Viral titers of VSVG-LV and BaEVRless-LV produced in parental HEK293T cells, as well as BaEVRless-LV from our optimized platform. Data are shown as mean ± standard deviation (SD) of three independent experiments (n=3). ns, not significant; *, p ≤ 0.05. (**D**) Microscopic images of parental HEK293T cells transfected with the *VSV-G* plasmid during lentiviral synthesis. (**E**) Images of HEK293T cells transfected with the *BaEVRless* plasmid under the same treatment. Scale bars are 500 μm in the 1X images and 100 μm in the zoomed images.

The pseudotype envelope protein can be modified to alter the lentiviral tropism by changing the envelope gene in the packaging plasmid. Upon expression, the envelope proteins localize to the cell membrane and are displayed on the surface of HEK293T cells. During virion release, a portion of the HEK293T cell membrane becomes the outer membrane of the lentivirus. For instance, the VSV-G envelope protein on the outer membrane of VSV-G-LV interacts with LDLR on the target cell, leading to the internalization of VSVG-LV and the release of lentiviral RNA transcript encoding the transgene. Similarly, the BaEVRless protein binds to ASCT1 and ASCT2 receptors on NK cells to facilitate viral entry.

We transduced NK cells with two types of lentiviruses encoding the *turbogfp* gene and assessed transduction efficiency by measuring turboGFP expression using flow cytometry. BaEVRless-LV demonstrated significantly higher efficiency than VSVG-LV in transducing primary human NK cells (Figure 1B). While the transduction efficiency of VSVG-LV plateaued at 20−40% even at high multiplicities of infection (MOI) of 50, BaEVRless-LV achieved an efficiency of 40−50% at an MOI of 1 and plateaued at approximately 80% at an MOI of 10. Our results confirm that BaEVRless-LV surpasses VSVG-LV in NK cell transduction efficiency across all MOI ranges.

However, BaEVRless-LV synthesis presents a unique challenge not observed in VSVG-LV. When packaged in HEK293T cells using the same protocol, the viral titer of BaEVRless-LV was approximately 100-fold lower than that of VSVG-LV (Figure 1C). Additionally, within 48 hours of transfection with the BaEVRless packaging plasmids, we observed extensive syncytia formation and the development of large sheets of fused cells (Figure 1D vs. 1E). These fused cells subsequently underwent cell death and detachment from the plate, resulting in the production of excessive cell debris and complicating lentivirus purification. Similar challenges have been reported previously (Girard-Gagnepain et al., 2014; Bauler et al., 2020; Noguchi et al., 2023).

### ASCT2 mediates BaEVRless-induced syncytia formation

We hypothesized that BaEVRless binds to the ASCT2 receptor on neighboring HEK293T cells, triggering membrane fusion (Figure 2A). This process may also involve the auxiliary receptor ASCT1 (Marin et al., 2000). To test this, we used CRISPR-Cas9 genome editing to KO the *ASCT1* and *ASCT2* genes. We designed four single-guide RNAs (sgRNA) targeting the exon 1 regions of *ASCT1* and *ASCT2* genes, and electroporated HEK293T cells with the four Cas9 RNPs independently. Three days post-electroporation, cells were analyzed to determine the percentage of insertion-deletion (% indel) and KO scores at the target site using Sanger sequencing (Figure 2B). ICE analysis revealed that Cas9 RNP 1 and 3 exhibited higher % indel and KO scores, indicating greater *ASCT1* and *ASCT2* KO efficiencies, respectively. Editing by Cas9 RNP 2 resulted in a nine-base in-frame deletion that did not disrupt ASCT1 expression, resulting in a low KO score.

**Figure 2.**
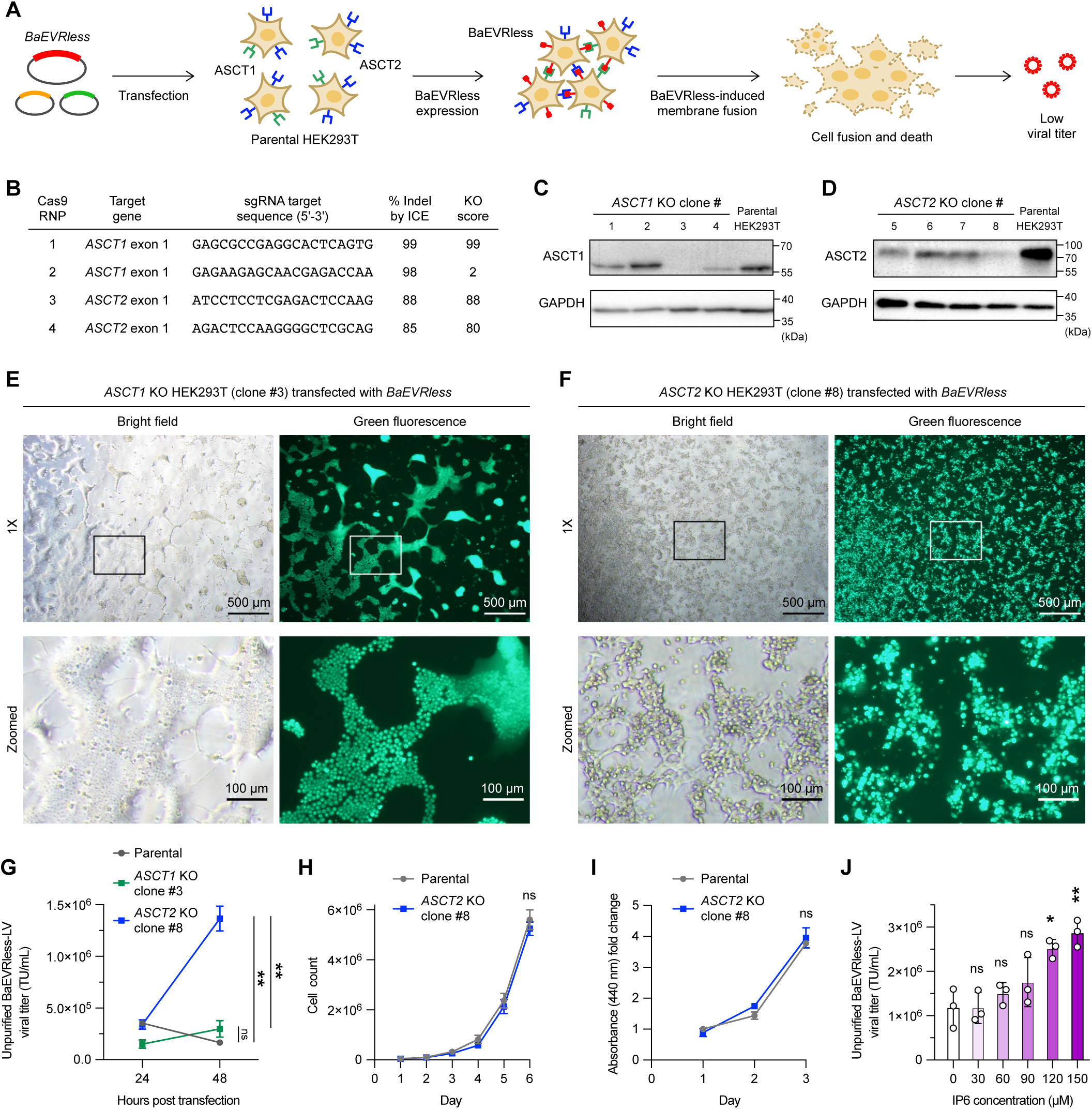
CRISPR knockout of the *ASCT2* gene alleviated BaEVRless-induced cell fusion. **(A)** Model of BaEVRless-induced cell fusion. BaEVRless engages the ASCT1 and ASCT2 receptors on neighboring HEK293T cells, leading to membrane fusion, cell death, and low BaEVRless-LV titer. **(B)** sgRNA sequences for CRISPR knockout (KO) of *ASCT1* and *ASCT2* genes in HEK293T cells. % insertion-deletion (indel) and KO scores were determined by Sanger sequencing and ICE analysis. **(C)** Immunoblotting of ASCT1 in four single *ASCT1*-KO clones versus parental HEK293T cells, with GAPDH as the loading control. **(D)** Immunoblotting of ASCT2 in four single *ASCT2*-KO clones. (**E**) Microscopic images of the *ASCT1*-KO clone #3 at 48 hours after transfection of *BaEVRless* plasmid. (**F**) Microscopic images of the *ASCT2*-KO clone #8 under the same treatment. Scale bars are 500 μm in the 1X images and 100 μm in the zoomed images. **(G)** Unpurified BaEVRless-LV titers from parental HEK293T, *ASCT1*-KO and *ASCT2*-KO cells at 24 and 48 hours post-transfection. (**H**) Cell proliferation rates of parental HEK293T and *ASCT2*-KO clone #8 under standard culture conditions, measured by cell counting. (**I**) Cell respiration rates of parental HEK293T and *ASCT2-*KO clone #8 under standard culture conditions, determined by WST-1 assay. **(J)** Unpurified BaEVRless-LV titers with increasing concentrations of inositol hexaphosphate (IP6) added during BaEVRless-LV synthesis. Data are shown as mean ± SD of three independent experiments (n=3). ns, not significant; *, p ≤ 0.05; **, p ≤ 0.01; ****, p ≤ 0.0001.

We performed fluorescence-activated cell sorting (FACS) of the Cas9 RNP 1- and 3-edited cells. We isolated four single clones for each KO and analyzed the ASCT1 or ASCT2 expression by immunoblotting. The majority of our KO clones showed a reduction in ASCT expression levels (Figure 2C and 2D). Specifically, *ASCT1* KO clone #3 and *ASCT2* KO clone #8 showed near ablation of ASCT1 and ASCT2 expression, respectively. Subsequently, we performed BaEVRless-LV packaging in clone #3 and #8, and monitored the cells 48 hours after BaEVRless-plasmid transfection. While syncytia formation was still observed in *ASCT1* KO clone #3, the effect was less pronounced compared to parental HEK293T (Figure 2E vs. Figure 1E). The *ASCT1* KO cells formed small patches of fused and adherent cell colonies, which eventually detached from the plate after 72 hours of incubation. In contrast, the *ASCT2* KO clone #8 exhibited a completely healthy cell morphology (Figure 2F). The *ASCT2* KO cells remained firmly attached to the plate and no syncytia were observed. Our findings pinpoint ASCT2 as the primary mediator of BaEVRless-induced syncytia, with ASCT1 likely playing an auxiliary role in this process.

### CRISPR KO of the *ASCT2* gene increases BaEVRless-LV titer

We wanted to determine whether preventing BaEVRless-induced syncytia could increase BaEVRless-LV synthesis. We performed BaEVRless-LV packaging in both parental and *ASCT2* KO cells, sampled the culture media at 24 and 48 hours, and measured the unpurified viral titers. Interestingly, we observed an eight-fold increase in viral titer in the *ASCT2* KO cells (Figure 2G), indicating that alleviating BaEVRless-induced syncytia increased BaEVRless-LV production. Furthermore, the *ASCT2*-KO cells showed normal proliferation and respiration rates as the parental HEK293T (Figure 2H and 2I). Since the *ASCT2*-KO cells could be maintained and efficiently transfected as the parental cells, these KO cells can be integrated seamlessly into the standard adherent cell-based lentiviral production system without needing modifications to the cell culture system or transfection procedures.

### Supplementation of inositol hexaphosphate increases BaEVRless-LV titer

Inositol hexaphosphate (IP6), also known as inositol hexakisphosphate or phytic acid, is a metabolite crucial for HIV-1 replication in human cells (Mallery et al., 2019). IP6 binds to the HIV-1 Gag protein and is packaged into the virion to stabilize the viral capsid. The cellular availability of IP6 directly impacts HIV-1 viral production and infectivity. Similarly, the Primate Lentivirus Simian Immunodeficiency Virus replies on IP6 for replication (Ricana et al., 2020). To assess if IP6 supplementation during BaEVRless-LV production could enhance viral titers, we added various concentrations of IP6 to *ASCT2*-KO cells during viral packaging. We observed a dose-dependent increase in BaEVRless-LV titer with IP6 supplementation (Figure 2J). At a concentration of 150 μM, IP6 boosted the viral titer by an additional three-fold. These findings suggest that IP6 supplementation, a non-toxic and naturally occurring metabolite, has the potential to enhance BaEVRless-LV production.

### Optimization of BaEVRless-LV purification and transduction methods are crucial

Solving the BaEVRless problem marks the initial step in a multi-step process in BaEVRless-LV transduction of primary NK cells. Optimization of viral purification and NK cell transduction protocols was also necessary. First, we compared two common lentivirus purification methods: sucrose-cushion centrifugation versus precipitation using Lenti-X concentrator, a commercial polyethylene glycol-based solution. We transfected the *ASCT2*-KO cells with BaEVRless-LV packaging plasmids and *turbogfp* transfer plasmid. BaEVRless-LV-containing media were collected after three days and subjected to the two purification methods (Figure S1A). We quantitated the viral titers before and after purification by transducing HEK293T cells and measuring turboGFP expression by flow cytometry. Sucrose-cushion centrifugation recovered 30% of the BaEVRless-LV from the unpurified media, whereas the Lenti-X precipitation allowed >90% recovery (Figure S1B). Notably, insoluble precipitation was observed after sucrose-cushion centrifugation, leading to a significant loss in viral yield. We also assessed NK cell viability post-transduction as an indicator of viral purity. Both purification methods showed similar cell viabilities in the range of 70−100%, suggesting comparable viral purities (Figure S1C). Ultimately, the combination of *ASCT2* KO cells, IP6 supplementation and an optimized purification protocol resulted in a 50-fold increase in BaEVRless-LV titer to approximately 5 × 10^8^ TU/mL, nearing the VSVG-LV titer (Figure 1C).

Next, we evaluated polybrene supplementation and spinfection in NK cell transduction using Lenti-X purified BaEVRless-LV encoding *turbogfp*. Primary NK cells from three donors were transduced, with the MOI reduced to 0.3 to sensitize the assay. Polybrene (hexadimethrine bromide), a cationic polymer, was added to enhance lentiviral transduction by neutralizing charge repulsion between virions and the cell surface (Denning et al., 2013). We tested three polybrene concentrations (2, 4, and 8 µg/mL) and observed a dosage-dependent increase in transduction efficiency, although high polybrene doses also reduced cell viability (Figure S1D and S1E). Spinfection, involving centrifugation of the NK cells and BaEVRless-LV mixture at 1200 × *g* for 90 min to enhance cell-virus contact, did not significantly improve transduction but compromised cell viability (Figure S1D and S1E). Consequently, we selected condition 3 (4 µg/mL of polybrene without spinfection) for all downstream experiments because of its optimal balance of transduction enhancement and cell viability.

### *ASCT2* KO cells enables robust packaging of large transgenes

Building on the *turbogfp* results, we proceeded to package two larger transgenes: the *EGFR-CAR* sequence and *TBX21* gene, encoding the T-bet transcription factor crucial for NK cell development (Fang et al., 2022). Co-expression of *turbogfp* or *egfp* with *EGFR-CAR* and *TBX21* simplified transduction analysis by flow cytometry (Figure 3A). BaEVRless-LV titers of *EGFR-CAR* (2147 nt) and *TBX21* (2400 nt) were comparable to that of *turbogfp* (717 nt), indicating robust packaging of large genes (Figure 3B). However, transduction and expression of *EGFR-CAR* and *TBX21* were less efficient compared to *turbogfp*. *TBX1* transduction efficiency was 30-50% at MOI 1 and increased to 60-80% at MOI 10 (Figure 3C). In contrast, *turbogfp* efficiency reached 70% at MOI 1 and plateaued at 80% at MOI 10 (Figure 1B). The result suggests that large transgenes may be less efficient to transduce or express in NK cells.

**Figure 3.**
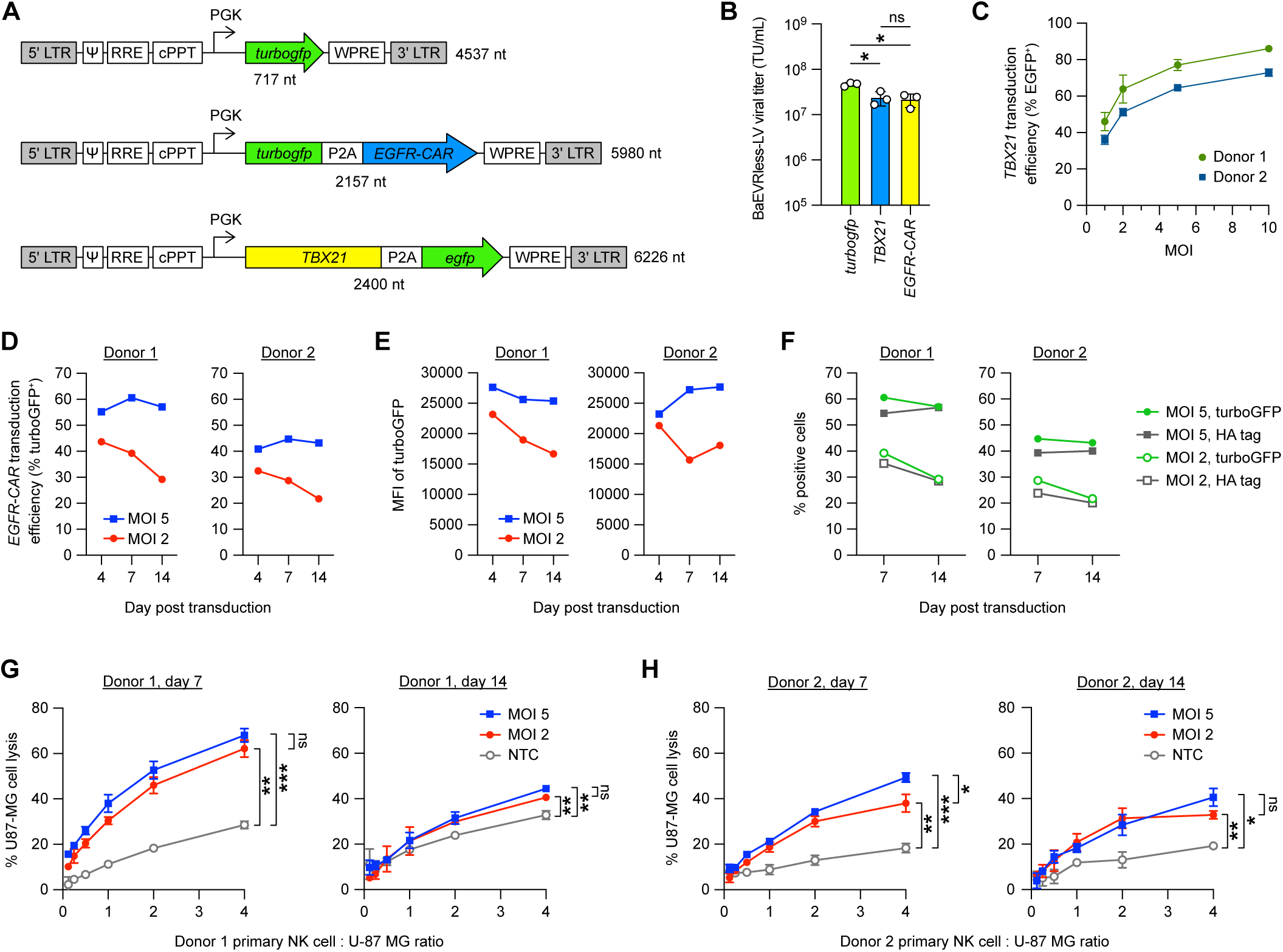
Robust packaging of large transgenes in BaEVRless-LV using the optimized synthesis platform. **(A)** Lentiviral transcripts encoding *turbogfp*, *TBX21* and EGFR-targeting chimeric antigen receptors (*EGFR-CAR*) sequences. The transgenes are expressed from the PGK promoter. The 5’ LTR, ψ, RRE, cPPT, WPRE and 3’ LTR are essential lentiviral elements. The *EGFR-CAR* and *TBX21* sequences are co-expressed with *turbogfp* and *egfp*, respectively, for measuring transduction efficiency by flow cytometry. **(B)** BaEVRless-LV titers encoding *EGFR-CAR* and *TBX21* were determined in HEK293T cells based on turboGFP and EGFP co-expression, respectively. **(C)** Transduction of *TBX21* in primary NK cells from two donors, with efficiencies estimated by EGFP co-expression at day 4, 7 and 14 post-transduction. (**D**) Transduction of *EGFR-CAR* in primary NK cells from two donors, with efficiencies estimated by turboGFP co-expression at day 4, 7 and 14 post-transduction. (**E**) Mean fluorescence intensity (MFI) of turboGFP. (**F**) Correlation of turboGFP and HA-tag expression levels in *EGFR-CAR*-transduced NK cells. (**G**) *In vitro* cytotoxicity of EGFR-CAR-expressing NK cells against U-87 MG glioblastoma cells compared to untreated NK cells (NTC). The assay was performed on day 7 and 14 post-transduction. (**H**) Cytotoxicity of donor 2 cells. Data are shown as mean ± SD of triplicate experiments (n=3). ns, not significant; *, p ≤ 0.05; **, p ≤ 0.01; ***, p ≤ 0.001.

A similar reduction in *EGFR-CAR* transduction was observed. We transduced two different donor cells at low MOIs of 2 and 5, and monitored the EGFR-CAR expression by turboGFP fluorescence on Day 4, 7 and 14 post-transduction. Transduction efficiency, measured by the % turboGFP^+^ cells, varied between the donor cells (Figure 3D). Also, the % turboGFP^+^ cells and fluorescence intensity of turboGFP declined over time (Figure 3E). To ensure turboGFP was a good indicator of EGFR-CAR expression, we also stained the HA affinity epitope within the CAR protein using an anti-HA antibody to detect EGFR-CAR protein displayed on NK cell surface. The percentages of HA^+^ and turboGFP^+^ cells were highly comparable in both donor cells at different MOIs, confirming the correlation of turboGFP and EGFR-CAR expression.

Next, we assessed the cytotoxicity of EGFR-CAR NK cells *in vitro* against U-87 MG, an EGFR^+^ glioblastoma cell line. EGFR-CAR NK cells showed higher cytotoxicity than the untransduced control NK cells (Figure 3G and 3H), indicating that EGFR-CAR was functional. The cytotoxicity of EGFR-CAR NK cells did not differ between MOI 2 and 5. Donor 1 CAR NK cells displayed high cytotoxicity on day 7, but the activity decreased significantly on day 14, likely due to reduced CAR expression. In contrast, the cytotoxicity of donor 2 CAR NK cells remained relatively consistent on days 7 and 14, suggesting donor variation in CAR-mediated cytotoxicity.

### Robust packaging of BaEVRless-LV enables CRISPR KO screen in primary NK cells

We expanded our study to include CRISPR KO screen in primary NK cells to verify the effectiveness of our BaEVRless-LV synthesis platform (Figure 4A). We adapted the SLICE editing protocol developed for human primary T cells (Shifrut et al., 2018), and repurposed it to screen for kinase genes crucial for primary NK cell proliferation. We selected the Brunello CRISPR kinome library, which comprises 6194 sgRNAs targeting 763 kinase genes in the human genome (Doench et al., 2016). The kinome sgRNA library was packaged in BaEVRless-LV using our optimized platform. To maintain a 1000-fold sgRNA coverage of the kinome library, we transduced 2 × 10^7^ NK cells with BaEVRless-LV at an MOI of 0.3, and electroporated the cells with Cas9 protein one day later. The pooled KO population was selected by puromycin treatment and then cultured for 10 days. Subsequently, we amplified the sgRNA cassette in the surviving NK cell population for next-generation sequencing (NGS) to identify genes essential for NK cell proliferation.

**Figure 4.**
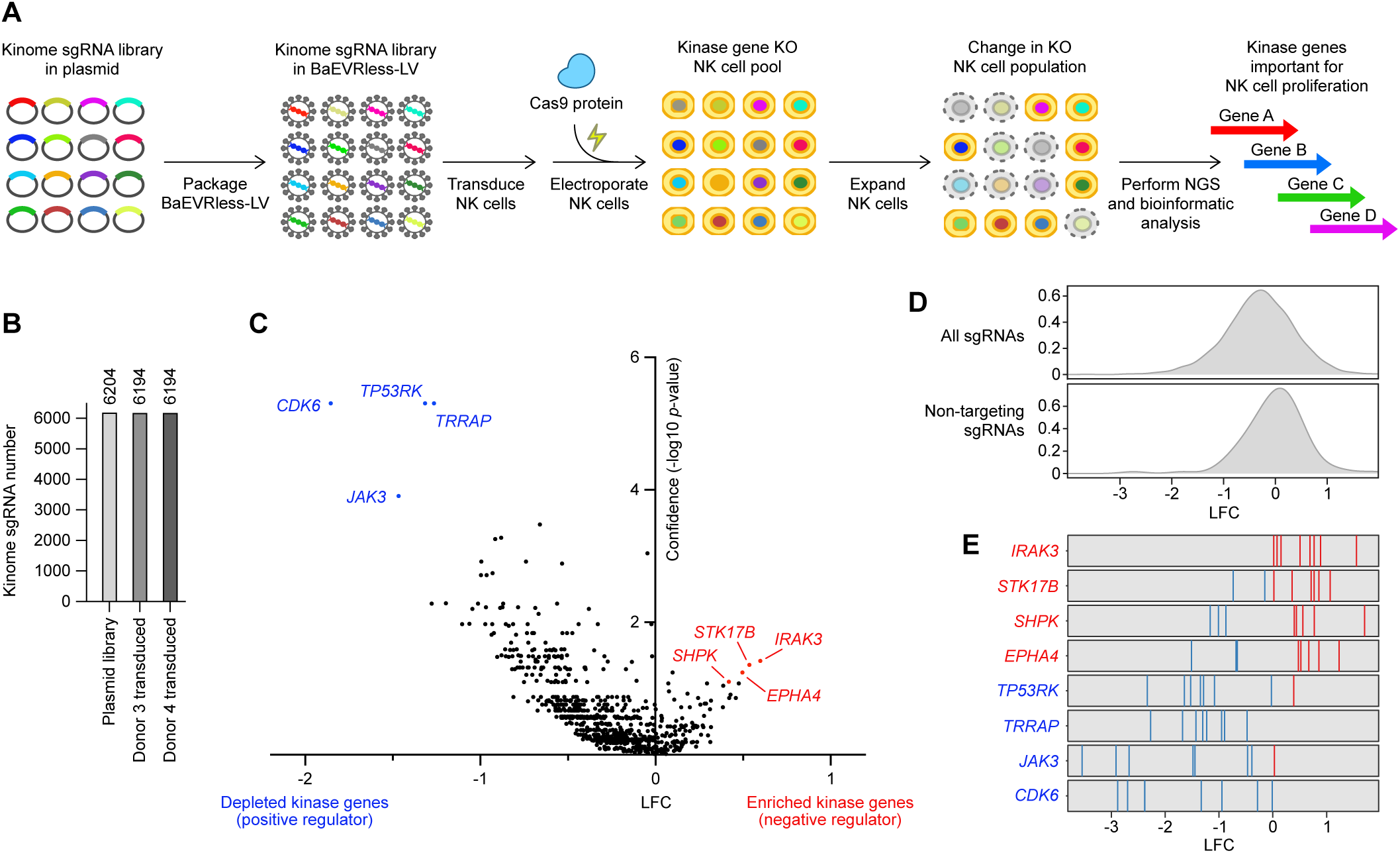
The kinome CRISPR KO screen in NK cells using BaEVRless-LV. (**A**) Workflow of the kinome CRISPR KO screen using the SLICE approach. The kinome CRISPR KO library, which contains 6204 sgRNAs targeting 763 kinase genes, was packaged in BaEVRless-LV for the transduction of primary NK cells. Cas9 protein was electroporated into NK cells to KO kinase genes, generating a population of unique KO cells. Change in KO cell distribution and corresponding sgRNA signature were determined by next-generation sequencing (NGS) and bioinformatic analysis. (**B**) Verification of the number of sgRNAs in the plasmid library and recovered from transduced NK cells by NGS. (**C**) Volcano plot showing the log2 fold-change (LFC) and confidence of enriched and depleted kinase genes from the kinome KO screen. All values were calculated from two donor cells as biological replicates. (**D**) Distribution of LFC values for 6094 targeting and 100 non-targeting sgRNAs in the kinome library. (**E**) LFC for all eight sgRNAs targeting four enriched genes (red lines) and four depleted genes (blue lines) in surviving NK cells.

The packaging and transduction of kinome sgRNA library were highly efficient in two independent experiments using two donor cells. After BaEVRless-LV transduction and puromycin selection, we analyzed the NK cells, and detected almost all sgRNAs in the Kinome library in both donor cells (Figure 4B). Depletion of sgRNAs targeting essential genes is an expected result of successful CRISPR KO screening (Bock et al., 2022). Indeed, we observed depletion of sgRNA targeting several important cell cycle kinase genes such as *CDK6* (Tigan et al., 2016), *TRRAP* (Tapias et al., 2014) as well as *TP53RK* (Peterson et al., 2010), a negative regulator of apoptosis response to mitotic stress (Figure 4C and 4D). We also observed depletion of *JAK3*, a downstream kinase of the IL-2 signaling pathway essential for NK cell survival (Robinette et al., 2018), indicating a successful screening outcome.

Some genes were enriched in our screen, albeit at lower fold-change, which is typical for CRISPR KO screens that are more effective at identifying depleted genes. *IRAK3* encodes IL-1 receptor–associated kinase-1, a key modulator of the innate inflammatory response that negatively regulates IL-1/TLR signaling (Nguyen et al., 2020). *IRAK3* KO in mice delayed cancer growth and enhanced the activation of myeloid and T cells (Tunalı et al., 2023). *STK17B* encodes the serine/threonine-protein kinase 17B, associated with T cell differentiation, activation and survival (Mandarano et al., 2020; Ryan and Craig, 2011). Phosphorylation of STK17B was upregulated in NK cells upon IL-2 and IL-15 stimulation (MacMullan et al., 2022). *EPHA4* encodes ephrin type-A receptor 4, which phosphorylates proline-rich tyrosine kinase 2, regulating the cytotoxic response between NK and target cells (Sancho et al., 2000; Aasheim et al., 2005). *SHPK* encodes sedoheptulose kinase of the pentose phosphate pathway fundamental to carbon homeostasis, nucleotide and amino acid biosynthesis, and oxidative stress response (Stincone et al., 2015). SHPK was reported to modulate glycolytic energy flux crucial to macrophage activation and polarization (Haschemi et al., 2012). While these enriched genes have known immune functions, their specific roles in NK cell immunity require further investigation.

In summary, our study elucidated the challenges posed by BaEVRless in lentiviral synthesis and addressed them through *ASCT2* KO, IP6 supplementation, and an optimized purification protocol. We demonstrated that BaEVRless-LV outperformed VSVG-LV in transducing primary human NK cells, despite initial hurdles in viral titer and syncytia formation. Through CRISPR-Cas9 genome editing, we identified ASCT2 as a key mediator of BaEVRless-induced syncytia, highlighting its role in BaEVRless-LV production. Furthermore, our study showcases the robust packaging capability of BaEVRless-LV for large transgenes and its utilities in CAR-NK manufacture and CRISPR KO screens in primary NK cells. The successful demonstration of the first CRISPR genetic screen in primary human NK cells, to our best knowledge, opens avenues for future genome-wide CRISPR screens and therapeutic developments. Our platform may also enable CRISPR screens in cells requiring BaEVRless-LV transduction such as primary B cells and HSPC. Overall, our work establishes BaEVRless-LV as a valuable tool for genetic manipulation in primary immune cells, with broad implications for cellular therapy and basic research.

## MATERIALS AND METHODS

### Reagents and antibodies

All chemicals were purchased from Merck and cell culture reagents from Thermo Fisher Scientific, unless specified otherwise. The following antibodies were used for immunoblotting: anti-GAPDH (#60004-1-Ig, ProteinTech), anti-ASCT1 (#8442, Cell Signaling Technology), anti-ASCT2 (#8057, Cell Signaling Technology), anti-rabbit IgG (#7074, Cell Signaling Technology) and anti-mouse IgG (#7076, Cell Signaling Technology). The following antibodies were used for flow cytometry analysis and purchased from BioLegend unless specified otherwise: APC anti-CD3 (#317317), PE anti-CD56 (#362507), PE anti-CD137L (also known as 4-1BB ligand, #311503), Alexa Fluor 647 anti-IL-21 (#513005), anti-HA epitope tag (#901513) and Alexa Fluor 647 anti-mouse IgG (Invitrogen #A21235).

### Cell culture

HEK293T, K562 and U-87 MG cell lines were purchased from the American Type Culture Collection (ATCC) and maintained according to the ATCC protocols. Briefly, HEK293T and U-87 MG were maintained in the Complete DMEM medium containing DMEM with high glucose (HyClone) supplemented with 10% heat-inactivated fetal bovine serum (FBS), 25 mM HEPES, 1X GlutaMAX, 1X penicillin-streptomycin (Pen-Strep). K562 was maintained in RPMI-1640 (ATCC modification) supplemented with 12.5% heat-inactivated FBS, 25 mM HEPES, 1X GlutaMAX, and 1X Pen-Strep. Genetically modified K562 was used as feeder cells to stimulate the *ex vivo* proliferation of primary NK cells. The modified K562 was generated by transduction with VSVG-LV expressing 4-1BBL and membrane-bound IL21 (mIL-21) (Denman et al., 2012). K562 feeder cells were expanded in RPMI medium containing 2 µg/ml of puromycin to maintain transgene expression and then irradiated at the dosage of 100 Gy to terminate cell proliferation. Irradiated K562 cells then were washed and used immediately for NK stimulation or cryopreserved at 3 × 10^6^ cells/vial in liquid nitrogen for future use. For routine passage, HEK293T and U-87 MG cells were dissociated by Trypsin-EDTA solution. The dissociated cells were diluted in fresh media at 1:4 to 1:8 ratio. Cell density and viability during routine passage were determined by Trypan blue staining in the Countess II cell counter (Thermo Fisher Scientific). All cell lines were maintained in a 37°C incubator with 5% CO_2_ and were routinely checked for Mycoplasma contamination using the EZ-PCR detection assay kit (Biological Industries).

### *ASCT* KO by Cas9 RNP electroporation

Cas9 protein and single-guide RNA (sgRNA) were prepared as described (Lin et al., 2022). The oligonucleotides for sgRNA template synthesis are listed in Table S1. *ASCT1*- and *ASCT2*-targeting sgRNAs were designed using CRISPR Design tool on Benchling website (www.benchling.com). Cas9 RNP electroporation was performed using Lonza 4D Nucleofector as described with modifications (Huang et al., 2021). Briefly, an electroporation reaction consisted of 2 × 10^5^ HEK293T cells in 20 μL of SF buffer (Lonza) and 2 μL of 20 μM Cas9 RNP (equivalent to 40 pmol final concentration). The cell mixture was then loaded into the 16-well nucleofection strips (Lonza) and electroporated by pulse code DS150. Immediately after electroporation, 100 μL of pre-warmed Complete RPMI 1640 medium was added to each well for cell recovery in a 37°C incubator for 15 minutes. Subsequently, the cells were transferred to 24-well culture plates filled with 1 mL of pre-warmed complete DMEM medium. Gene-editing analyses were conducted 72 hours post-electroporation.

### *ASCT* KO analysis by Sanger sequencing and ICE

*ASCT* KO efficiency was determined by Sanger sequencing and ICE analysis using the default parameters (https://ice.synthego.com). Briefly, the edited HEK293T cells were collected by trypsin dissociation, pelleted by centrifugation at 300 × g for 5 min and washed with DPBS once. Cell pellets were lysed by QuickExtraction solution (Lucigen) per the manufacturer’s instruction at 65°C for 15 min, 98°C for 5 min and 4°C for 10 min to extract the genomic DNA. One hundred ng of genomic DNA was used for PCR amplification of the target loci using KAPA HiFi HotStart PCR kit (Roche) and the primer sets in Table S1. After validated by DNA gel electrophoresis, the PCR product was purified by QIAquick PCR Purification Kit (Qiagen), eluted in H_2_O and subjected to Sanger sequencing at the Institute of Biomedical Sciences, Academia Sinica. The % indel was determined on the Synthego website by Inference of CRISPR Edits tool (ICE) using the default setting (https://www.synthego.com/products/bioinformatics/crispr-analysis).

### Immunoblotting

ASCT1 and ASCT2 expression were detected by immunoblotting. The KO HEK293T cell pellet was dissolved in 50 μL of SDS loading dye containing 80 mM Tris (pH 6.8), 2% β-mercaptoethanol, 4% SDS, 15% glycerol and 0.01% Orange G. The cell lysate was heated at 95°C for 5 mins, and resolved by electrophoresis in 10% SDS-PAGE gel at 100 V for 1.5 hour. The proteins were transferred from SDS-PAGE to PVDF membrane (0.2-μm, Millipore) in Tris-Glycine Transfer buffer (Bio-Rad) by Trans-Blot SD Semi-Dry Transfer (Bio-Rad) at 80 mA for 1.5 hours. The membrane was incubated in TBST with 5% (w/v) skim milk (BD Biosciences) for 1 hour at room temperature. After blocking, the membrane was incubated with the primary antibodies in TBST with 5% skim milk overnight at 4°C. The membrane was washed three times with TBST and incubated with the secondary antibodies in TBST with 0.5% BSA for 1 hour at room temperature. The membrane was washed three times with TBST. The target proteins were visualized by Western Lightning PLUS-ECL or Ultra kit (Perkin Elmer). The images were acquired by Azure Biosystems C300 or Thermo Fisher Scientific iBright FL1000, and processed by ImageJ.

### Flow cytometry and FACS

Flow cytometry was performed on CytoFLEX (Beckman Coulter) at the Flow cytometry core facility in the Institute of Biological Chemistry, Academia Sinica. NK cells were harvested by centrifugation at 500 × *g* for 5 min and washed once with 1 mL of ice-cold Flow buffer (DPBS supplemented with 2% FBS, 25 mM HEPES, and 0.5 mM EDTA). For surface protein detection, the cells were stained with antibody solution at the manufacturer’s recommended ratio in the dark on ice for 15 min. After staining, the cells were washed with 1 mL of Flow buffer, pelleted at 500 × *g* for 5 min, resuspended in 200 μL of Flow buffer, and transferred to a 5-mL Falcon polystyrene flow tube with a cell strainer snap cap (Corning). Modified K562 feeder cells were prepared similarly to monitor the expression the 4-1BBL and mIL-21. The *ASCT1*- and *ASCT2*-KO HEK293T cells were single cell-sorted by FACSAria III (BD Biosciences) at the flow cytometry core facility in the Institute of Biomedical Sciences, Academia Sinica. The isolated cells were collected in 96-well plates at one cell per well in 200 μL of the Complete DMEM medium and expanded by the standard culture method. The expanded cells were validated for the ablation of ASCT1 and ASCT2 protein expression by immunoblotting before storing in liquid nitrogen. All data were analyzed using FlowJo (BD Biosciences) and CytExpert (Beckman Coulter) software.

### Construction of BaEVRless envelope plasmid

Packaging plasmid pCMV.deltaR8.91 and VSV-G envelope plasmid pMD2.G were obtained from C6 RNAiCore facility in Academia Sinica. BaEVRless envelope plasmid was generated by replacing the *VSV-G* gene in pMD2.G plasmid with the BaEVRless sequence reported previously (Girard-Gagnepain et al., 2014). The PCR primers are listed in Table S2. Briefly, the BaEVRless gene was codon-optimized and synthesized as a gBlock fragment from IDT DNA. The BaEVRless sequence was amplified from the gBlock by KAPA HiFi HotStart PCR kit using primer 1 and 2. The pMD2.G vector backbone without the *VSV-G* gene was amplified by primer 3 and 4. The PCR products were resolved by DNA gel electrophoresis, extracted by QIAquick PCR Purification Kit, and ligated by NEBuilder HiFi DNA Assembly (NEB) per the manufacturer’s protocol. The plasmid was transformed and maintained in *E. coli* Stbl3 strain (Thermo Fisher Scientific) and purified using Qiagen plasmid extraction kit. BaEVRless plasmid was validated by Sanger Sequencing in DNA Sequencing Core Facility at the Institute of Biomedical Sciences at Academia Sinica. BaEVRless plasmid map (pMD2.G_BaEVRless) is attached as Genbank file in the Supplementary Data.

### Construction of lentiviral transfer plasmids

Lentiviral transfer plasmids encoding different transgenes were constructed into the pHR vector (modified from Addgene #79125) by NEBuilder HiFi DNA Assembly (NEB) using the same approach as BaEVRless plasmid. The PCR reactions were performed using KAPA HiFi HotStart PCR kit and the primers are listed in Table S2. Briefly, to make the pHR_turboGFP plasmid, the *turbogfp* gene was amplified from pmaxGFP plasmid (Lonza) using primer 5 and 6, and ligated with the pHR vector amplified by primer 7 and 8. To make the pHR_TBX21_P2A_EGFP plasmid, the *TBX21* gene was amplified from the cDNA of NK cells using primer 9 and 10, and ligated with the pHR vector amplified by primer 11 and 12. The EGFR-CAR expression cassette contains the following sequences: *turbogfp* gene, T2A, CSF2RA signal peptide, anti-EGFR single-chain variable fragment (scFv) from Panitumumab, IgG hinge, CD8 transmembrane domain, 4-1BB co-stimulating domain and CD3ζ activation domain. The CAR cassette was synthesized as a gBlock fragment from IDT DNA, amplified by primer 13 and 14, and ligated into pHR vector amplified by primer 15 and 16. All plasmids were validated by Sanger sequencing. The plasmid maps are attached as Genbank files in the Supplementary Data.

### Lentiviral production

HEK293T and the *ASCT* KO cells were maintained as described above. The cells were passaged every 2−3 days at the confluency of 80-90%. To ensure consistent and high lentiviral titer, it is recommended not to use cells that have been passaged more than 30 times. On day 0, 8.5−9.0 × 10^6^ cells were seeded on a 10-cm dish in 10 mL of lentiviral packaging medium (Opti-MEM medium supplemented with 5% heat-inactivated FBS, 1X GlutaMAX, 1X sodium pyruvate and 1X MEM non-essential amino acids solution). Approximately 16-18 hours after seeding, the cells were transfected with 8 µg of transfer plasmid, 7 µg of packaging plasmid (pCMV deltaR8.91), 3 µg of envelope plasmid (pMD2.G or pMD2.G_BaEVRless) using Lipofectamine 3000 according to the manufacturer’s instruction. The medium was removed 6 hours after transfection, and replenished with fresh lentiviral packaging medium supplemented with IP6 (also known as phytic acid). The 30 mM stock IP6 solution was prepared by dissolving IP6 powder (Merck) in DPBS, pH-adjusted to 7, filter-sterilized and stored at 4℃. IP6 was added at the concentrations indicated in Figure 2J or at the optimal concentration of 150 µM. At 48-50 hours post transfection, the lentivirus-containing medium was collected in a 15-mL conical tube and centrifuged at 650 × *g* for 5 minutes to remove cell debris. The supernatant was passed through low-protein binding 0.45-µm syringe filter (Millipore). The lentivirus was purified by two different methods for comparison. In the first method, lentivirus was precipitated from the supernatant using Lenti-X Concentrator (Takara) according to the manufacturer’s protocol. In the second method, lentivirus was pelleted by centrifugation in 20% sucrose cushion at 4500 × *g* at 4℃ for 20 hours. In both methods, the lentivirus pellet was suspended at 1/50 volume of the cell culture supernatant in the basal NK MACS medium or Opti-MEM. The purified lentivirus was snap-frozen in liquid nitrogen as aliquots and store at −80°C.

The lentiviral vectors in this study all carry the *gfp* reporter gene for detection, except the kinome CRISPR sgRNA library. The functional titer of lentivirus was determined by transducing HEK293T cells and measuring transduction efficiency by flow cytometry based on GFP expression. Briefly, 5 × 10^4^ HEK293T cells were seeded in 500 µL of Complete DMEM medium containing 8 µg/mL of polybrene in 24-well plate. Different amount of lentivirus ranging from 0.1 to 5 µL were added to transduce HEK293T cells for 2 days. Gene expression was determined by flow cytometry and functional titer was calculated based on the samples that gave rise to 10−30% transduction efficiency following the formulation below:

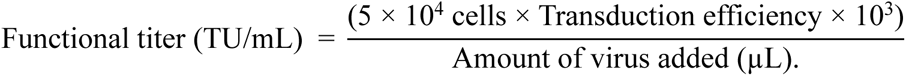

Functional titer (TU/mL) = (5 × 10^4^ cells × Transduction efficiency × 10^3^) Amount of virus added (µL).

The lentiviral titer of CRISPR sgRNA library was determined based on puromycin resistance. HEK293T cells were transduced with different amounts of lentivirus and treated with 2 µg/mL of puromycin on day 2 after transduction. On day 4, 100% of non-transduced cells should be killed by puromycin. Transduction efficiency was determined based on the number of live cells in the transduced group by Trypan Blue staining compared to the cell number in non-transduced group without puromycin treatment. Functional titer was calculated from the samples that gave rise to 30% transduction efficiency.

### Imaging analysis of BaEVRless-induced syncytia

The parental, *ASCT1* KO (clone #3) and *ASCT2* KO (clone #8) HEK293T cells were transfected by lipofectamine with lentiviral packaging plasmids encoding the *VSV-G* or *BaEVRless* envelope gene and a transfer plasmid encoding the *turbogfp* gene as described above. No IP6 was added. At 48 hours post transfection, bright-field and green fluorescent images of the parental cells and KO clones were taken on an Olympus CKX41 inverted microscope and processed by ImageJ version 1.53A.

### *Ex vivo* expansion of primary NK cells

CD3 negatively enriched and cryopreserved human peripheral blood NK cells were purchased from Lonza and HemaCare. Donor information is summarized in Table S3. All experiments on human cells were conducted according to the human experimental guidelines approved by the Institutional Review Board on Biomedical Science Research, Academia Sinica. On day 0, the cryopreserved NK cells were thawed in a 37°C water bath with gentle agitation until no ice was observed inside the tube. NK cells were gently transferred to a 15-mL conical tube (Falcon) containing 8 mL of 4°C RPMI 1640 (ATCC modification) medium supplemented with 10 U/mL DNaseI (Merck). The cryovial was rinsed with 1 mL of the RPMI medium to transfer the remaining NK cells to the 15-mL tube. After centrifugation at 400 × *g* for 10 min, the cell pellet was resuspended in 10 mL of prewarmed Complete NK MACS medium containing NK MACS Medium (Miltenyi), 5% EliteGro-Adv human plate lysate (EliteCell), 1% GlutaMAX, and 1% Pen-Strep. The medium was also supplemented with 10 U/ml of DNaseI and 2 ng/mL of recombinant IL-15 (Peprotech). The cell suspension was transferred to a T25 flask (Corning) and maintained in a 37°C incubator with 5% CO_2_. On day 1, NK cells were collected from the flask and the cell density was determined to set up the feeder-dependent expansion. Approximately 5 × 10^6^ NK cells were co-cultured with 1 × 10^7^ of the modified and irradiated K562 feeder cells in 40 mL of Complete NK MACS medium supplemented with 100 U/mL of recombinant IL-2 (Peprotech) in a T75 flask (Corning). The final NK cell density was 1.25 × 10^5^ cells/mL in 40 mL of medium. On day 3 and 5, NK cells were pelleted by centrifugation at 300 × *g* for 5 minutes. Twenty mL of the supernatant was carefully removed from the top layer and replaced with 20 mL of fresh Complete NK MACS medium with 100 U/mL IL-2. NK cells were gently resuspended and transferred back to the flask. On day 7, NK cells were collected by centrifugation at 300 × *g* for 5 minutes, washed once with DPBS (Corning) and cryopreserved in CryoStor (Sigma Aldrich) at 3 × 10^6^ NK cells per vial for future use. The remaining NK cells are re-stimulated with K562 feeder cells at ratio 1:1 and maintained by the same method above. The primary NK cells were maintained at a concentration below 3 × 10^6^ cells/mL and were passaged every 2−3 days by dilution in fresh Complete NK MACS medium. The purity of primary NK cells was analyzed by flow cytometry based on CD3 and CD56 expression.

### NK cell transduction

After 7−10 days of expansion, 5 × 10^4^ of primary NK cells were suspended in 100 µL of fresh Complete NK MACS medium per well of the 96-well flat-bottom plate. Lentivirus was added at the MOI indicated in the experiment. Polybrene, also known as hexadimethrine bromide, was purchased from Merck, dissolved in H_2_O to 8 mg/mL and sterilized by passing through a 0.2-µm filter. Polybrene solution was stored at -80°C as aliquots to prevent repeated freezing and thawing. Polybrene was supplemented in the medium at concentrations as specified in Figure S1D or at 4 µg/ml for all other experiments. Spinfection was performed at 1200 × *g* for 90 min at 32°C. Afterward, the cells were resuspended by gentle pipetting and incubated at 37°C overnight. One day after transduction, the cells were washed and resuspended in 100 µL fresh Complete NK MACS medium. The transduced cells were analyzed by flow cytometry to determine the transduction efficiency at the specified time.

### Respiration assay

Cellular respiration was determined by WST-1 assay (Abcam) using the manufacturer’s protocol. Briefly, parental and the *ASCT2* KO HEK293T cells were seeded in a 12-well plate at 1 × 10^4^ cells in 1 mL of Complete DMEM medium per well, and set up three wells per cell type. After one day of incubation, 100 µL of the medium was removed from each well, and 100 µL of WST-1 reagent was gently added to the cell culture to avoid disturbance of the cells. The plate was gently rocked to evenly mix WST-1 reagent with the culture medium, and then incubated at 37°C for 2 hours. After incubation, 100 µL of culture supernatant was transferred to a transparent 96-well plate and the absorbance at 440 nm and 650 nm (reference) was measured in Infinite M1000 Pro microplate reader (Tecan).

### CAR-NK cell cytotoxicity assay

Calcein-AM-based cytotoxicity assay was as described by Huang et al., 2021. U-87 MG cells were dissociated by trypsin (Gibco), neutralized by culture medium and pelleted at 200 × *g* for 3 min. The cells were wash once with DPBS, and adjusted to 1 × 10^6^ cells in 1 mL of DPBS containing 10 µM of Calcein-AM (BioLegend) for cell staining at 37°C for 30 min. Next, the cells were washed three times with DPBS and resuspended to 1 × 10^5^ cells/mL in RPMI-1640 (ATCC modification). CAR-NK cells were pelleted at 90 × *g* for 10 min and resuspended to 4 × 10^5^ cells/mL in RPMI-1640 (ATCC modification). One hundred µL of CAR-NK cells per well were added to a round-bottom 96-well plate. Serial dilution was performed for different ratios of CAR-NK to U-87 MG cells. One hundred µL of the stained U-87 MG cell suspension was added into each well to start the cytotoxicity assay. The 96-well plate was centrifuged at 120 × *g* for 3 min to initiate the contact between NK and U-87 MG cells. The cell mixture was incubated at 37°C for 4 hours. Spontaneous release of Calcein-AM by U-87 MG cells was measured in the absence of CAR-NK cells. Maximal release was determined by complete lysis of U-87 MG cells in RPMI-1640 (ATCC modification) containing 2% Triton-X100. After co-culture, the plates were centrifuged at 120 × *g* for 3 min, and 100 µL of the supernatant was transferred to 96-well Opti-plates (PerkinElmer). The 488/520 values were recorded by Infinite M1000 pro (Tecan). The following equation was used for cytotoxicity calculation:

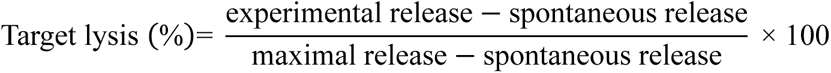

### The kinome CRISPR knockout screen using the SLICE approach

The kinome CRISPR knockout screen was performed in primary NK cells using the SLICE (sgRNA lentiviral infection with Cas9 protein electroporation) approach developed for primary T cells (Shifrut et al., 2018). Human Kinome CRISPR Knockout Library was from Addgene (catalog #1000000082) and prepared as described (Doench et al., 2016). The kinome CRISPR sgRNA library was packaged into BaEVRless-LV in the *ASCT2* KO cells using our standard protocol. The viral titer was determined in HEK293T cells by puromycin selection as described above. About 1 × 10^7^ of the feeder-expanded NK cells were transduced with the BaEVRless-LV at MOI 0.3. Recombinant Cas9 protein was prepared as detailed in (Lin et al., 2022) At 24 hours after transduction, Cas9 was electroporated to the NK cells using Lonza 4D nucleofection system using pulse code CM137 in 100-µL electroporation cuvettes. The reaction contained 400 pmol of Cas9 protein and 1 × 10^7^ of the transduced NK cells in 100 µL of P3 solution (Lonza). Immediately after electroporation, 500 µL of prewarmed Complete NK MACS medium was added directly into the cuvette to recover the NK cells for 15 min at 37°C. Next, NK cells were transferred to a T25 flask at 1 × 10^6^ cells/mL, and incubated at 37°C incubator. After 24 hours, the NK cells were collected by centrifugation at 300 × *g* for 5 min, and resuspended in the Complete NK MACS medium supplemented with 1 µg/mL of puromycin to select for the sgRNA-transduced cells selected for 48 hours. Next, the NK cells were collected by centrifugation at 300 × *g* for 5 min, washed once with DPBS, and resuspended in the puromycin-free Complete NK MACS medium. The NK cells were cultured for 7 days after puromycin selection, during which fresh medium was added every 2–3 days to maintain the NK cell density at 5 × 10^5^ to 2 × 10^6^ cells per mL. The cell suspension was split into multiple flasks when needed. The total cell number was maintained at 3 × 10^6^ cells minimum to preserve complete sgRNA representation. On day 21, we harvested 3 × 10^6^ cells (equivalent to 1000X coverage) for genomic DNA extraction and NGS to determine the sgRNA sequence in the NK cell population.

### Preparation of genomic DNA for NGS

The genomic DNA of NK cells were purified using Quick DNA Miniprep Plus kit (Zymo Research) according to manufacturer’s protocol. Amplification and bar-coding of the sgRNA sequences from genomic DNA were performed using a two-step PCR reaction as described (Shifrut et al., 2018). The PCR primers are listed in Table S4. The reaction consisted of 50 µL of NEBNext High Fidelity 2X PCR Master Mix (NEB), 4 µg of genomic DNA, 2.5 µL each of the 10 µM NG-Lib-Fwd and NGS-Lib-KO-Rev primers, and water to 100 µL total. The thermocycler setting was: 3 minutes at 98°C, followed by 10 seconds at 98°C, 10 seconds at 66°C, 25 seconds at 72°C, for 23 cycles; and a final 2 minutes extension at 72°C. The PCR amplicons were purified by DNA gel electrophoresis and extracted using Zymoclean Gel DNA Recovery kit (Zymo Research). The concentration and size of the amplicons were determined by Qubit DNA quantification (Thermo Fisher Scientific) and Fragment Analyzer (Agilent). The Nextera XT Index Kit v2 (Illumina) was used to add the dual-barcoded adaptors. The molar concentration of the amplicon from each sample was normalized and then sequenced on a NextSeq 500/550 instrument (Illumina). The purified Kinome CRISPR Knockout Library plasmids were also PCR-amplified and sequenced using the same procedure to verify the integrity of the sgRNA library.

### Analysis of pooled CRISPR Screens

The enriched and depleted genes in the kinome screen were identified using the MAGeCK software (Li et al., 2014). The read count table of each sgRNA was determined from raw fastq sequencing files by the default MaGeCK “count” module. The “–trim-5 48,49,50,51,52” parameter was used to retrieve sgRNA with offset design during the count table construction. To obtain gene level enrichment, the MAGeCK “test” module was performed with default settings using the alpha-robust rank aggregation (RRA) algorithm. The non-targeting control sgRNA in each library was used to calculate the size factor for normalization between each experiment. The distribution of individual sgRNA from the top-ranking genes is visualized using the MAGeCKFlute R package with default settings. The gene list from MAGeCK analysis is in Table S5. The original NGS data files are available at the NCBI Sequence Read Archive (PRJNA1078135).

### Statistical analyses

Except for the screening experiments, all data were collected from three independent experiments to determine mean values ± standard deviation as shown. Unpaired Welch’s unequal variances *t*-test was used to test for significant differences between two groups. P-values ≤ 0.05 were considered statistically significant. Statistical analyses were performed using GraphPad Prism 9.

## Figure and table legends

**Figure S1.**
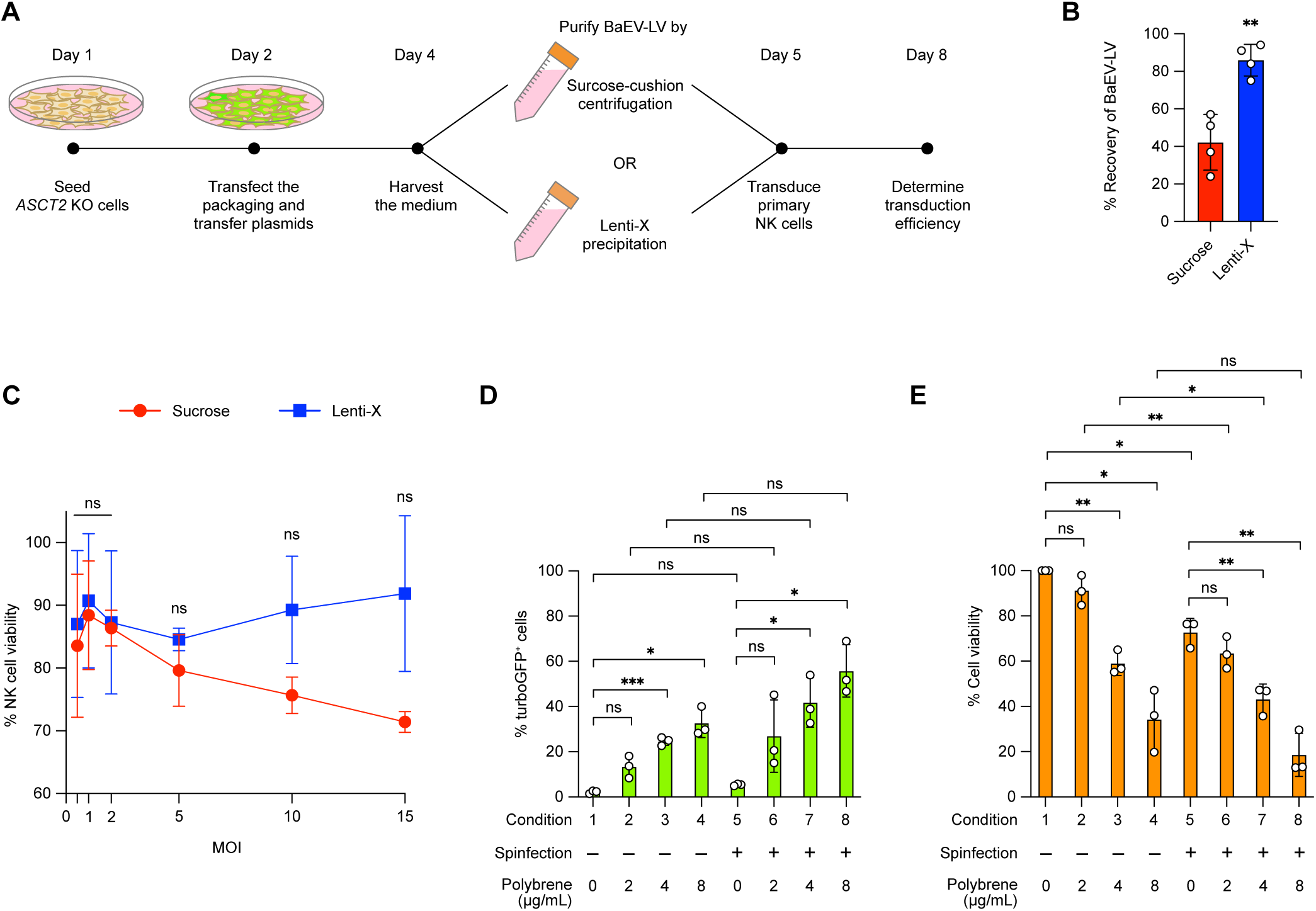
Optimization of BaEVRless-LV purification and NK cell transduction. (**A**) Workflow of BaEVRless-LV purification by sucrose-cushion centrifugation or Lenti-X precipitation. (**B**) Recovery rates of BaEVRless-LV by the two purification methods. Data are shown as mean ± SD of four independent experiments (n=4). (**C**) NK cell viability at 72 hours after transduction by BaEVRless-LV from the two purification methods. (**D**) Transduction efficiencies of BaEVRless-LV, encoding the *turbogfp* gene, at MOI 0.3 under different spinfection and polybrene treatments. (**E**) The viabilities of BaEVRless-LV-transduced NK cells under different spinfection and polybrene treatments. Data in C, D and E are shown as mean ± SD of three donor cells (n=3). ns, not significant; *, p ≤ 0.05; **, p ≤ 0.01; ***, p ≤ 0.001.

**Table S1: Oligonucleotides for sgRNA template synthesis and ICE analysis.**

**Table S2: Primer list for plasmid construction.**

**Table S3: Donor information of the cryopreserved primary NK cells.**

**Table S4: Primer list for NGS.**

**Table S5: Gene hits from kinome CRISPR KO screen generated by MaGeCK.**

## Acknowledgments

We thank members of the Lin lab for insightful discussions. We thank the IMB Genomics Core Facility at the Institute of Molecular Biology in Academia Sinica for conducting the NGS library preparation and sequencing. This work was supported by the National Science and Technology Council (https://www.nstc.gov.tw, grant number MOST 111-2311-B-001-017) and Academia Sinica (https://www.sinica.edu.tw). The funders had no role in the study design, data collection and analysis, decision to publish, or preparation of the manuscript.

## Author contributions

Y.J.L., Q.V.N. and S.L. conceived and designed this study. Y.J.L., Q.V.N., T.L.C. and K.L.Y. performed the experiments. Y.J.L. analyzed the NGS results. Y.J.L., Q.V.N. and S.L. wrote the manuscript.

## Disclosures

The Authors declare no competing financial interest.

## Notes

### Competing Interest Statement

The authors have declared no competing interest.

